# A 3D molecular map of the cavefish neural plate illuminates eyefield organization and its borders in vertebrates

**DOI:** 10.1101/2021.05.05.442716

**Authors:** François Agnès, Jorge Torres-Paz, Pauline Michel, Sylvie Rétaux

**Affiliations:** Institut des Neurosciences Paris-Saclay, Université Paris-Saclay, CNRS UMR9197, 91190Gif-sur-Yvette, France

**Author notes:** Equal contribution to this work (François Agnès, Jorge Torres-Paz).

**Keywords:** single cell, cell identity, eye transcription factors, telencephalon, hypothalamus, *Astyanax mexicanus*, natural variation, fluorescent ISH

## Abstract

The vertebrate retinas originate from a specific anlage in the anterior neural plate called the eyefield. The eyefield shares its anterior border with the prospective telencephalon and is in contact ventrally and posteriorly with hypothalamic and diencephalic precursors. Eyefield identity is conferred by a set of “eye transcription factors”, whose combinatorial expression has not been precisely characterized. Here, we use the dimorphic teleost species *Astyanax mexicanus*, which develops proper eyes in the wild type and smaller colobomatous eyes in the blind cavefish embryo, to unravel the molecular anatomy of the eyefield and its micro-evolutionary variations in the two *Astyanax* morphs. Using a series of markers (*Rx3, Pax6, CxCr4b, Zic1, Lhx2, Emx3, Nkx2.1*), we draw a comparative 3D expression map at the end of gastrulation/onset of neurulation, which highlights hyper-regionalization of the eyefield into sub-territories of distinct sizes, shapes, cell identities and putative fates along the three body axes. We also discover sub-domains within the prospective telencephalon, and we characterize cell identities at the frontiers of the eyefield. Analyses at the tissue scale and single cell level show variations in volumes and shapes of eyefield subdivisions as well as cellular gene expression levels and identity changes in cavefish. The ventro-medial border and the anterior border of the eyefield contain cells co-expressing hypothalamic and telencephalic marker, respectively, in cavefish embryos. Altogether, we provide a new model of eyefield patterning in 3D and suggest a developmental origin for the emergence of the coloboma phenotype in the natural mutant cavefish embryo.

## Introduction

In vertebrates, the eye retinas emerge from the anterior neural plate at the end of gastrulation, from a presumptive territory called the eyefield. Fate maps in mouse (Inoue et al., 2000), frog (Eagleson and Harris, 1990), bird (Cobos et al., 2001) or fish embryos (Varga et al., 1999; Woo and Fraser, 1995) have shown that the eyefield occupies a central position in the neural plate, between the telencephalon anteriorly and the diencephalon posteriorly. At such early stage of eye development, the eyefield already displays significant variations of size and shape across species, which likely prefigure their future morpho-anatomical differences, and which probably result from species-specific fine-tuning of earlier inductive and signalling events (Bielen et al., 2017; Rétaux et al., 2013).

During gastrulation, anterior neural fate acquisition and progressive distinction between telencephalon or eyefield identities require antagonizing posterior Wnt signals by secreting anti-Wnts from the anterior neural border (Heisenberg et al., 2001; Houart et al., 2002; Houart et al., 1998), as well as restricting the Bmp pathway activity in the anterior-most neural ectoderm (Bielen and Houart, 2012). The later allows protecting the future telencephalon from acquiring eye identity by repressing the expression of the key eyefield transcription factor rx3 (Bielen and Houart, 2012; Fish et al., 2004; Stigloher et al., 2006; reviewed in Giger and Houart, 2018). In fact, the eyefield transcription factors *rx, pax6, six3/six6 and lhx2*, detectable prior to neurulation, are expressed in dynamic, overlapping patterns and constitute a genetic network regulating vertebrate eye development (Zuber et al., 2003). A model for progressive eyefield specification was proposed, in which *otx2* initially primes the anterior neural plate for eyefield formation, leading eyefield transcription factors to form a self-regulating feedback network. Eyefield identity thus relies on the combinatorial expression of a limited number of factors, which define together a single medial domain, before its rapid evagination in two symmetric optic vesicles.

During neurulation, the eyefield program initiates eye morphogenesis and segregates eye-fated cells from other forebrain cells. Hence, anterior neural plate cells adopt specific and drastically different migratory behaviours and trajectories according to their identity, i.e dorso-medial convergence for telencephalic progenitors, evagination for eyefield cells and anterior ward movement for hypothalamic precursors (reviewed in Bazin-Lopez et al., 2015; Sinn and Wittbrodt, 2013; Wilson and Houart, 2004). Among them, eye progenitors follow complex movements, starting even before the onset of evagination of the bilateral optic vesicles (England et al., 2006). Live imaging also revealed that subdomains of the optic vesicles do maintain their relative organizations during morphogenesis (Kwan et al., 2012), suggesting that related, adjacent retina territories might be already determined at late gastrula before evagination.

Anteriorly at neural plate stage, the eyefield juxtaposes the prospective telencephalon, as revealed by specific markers in zebrafish (Stigloher et al., 2006). By contrast, eyefield posterior limits are less clear, as suggested by *pax6* extending posteriorly to *rx3* (Loosli et al., 2001; Zuber et al., 2003) and overlapping with diencephalic markers (Macdonald et al., 1997; Staudt and Houart, 2007). Thus, the master eye gene *pax6* (Ashery-Padan and Gruss, 2001) and *rx3*, which is also necessary and sufficient for eye formation in fish (Ishikawa et al., 2001; Loosli et al., 2003), do not appear to form a uniform optic domain. Further, the other eyefield transcription factors must add supplementary combinatorial patterning complexity to the eyefield, which have yet to be investigated.

Here, we undertook a detailed molecular portrait of the eyefield, using several markers to define cell identities within this anterior neural plate region and to determine its exact frontiers with surrounding territories in the 3D. For this purpose, we used the fish *Astyanax mexicanus* embryos as a model system. Through its intra-species dimorphism represented by wildtype river-dwelling and blind cave-dwelling morphotypes, it offers an exquisite model to unravel subtle variations in body plans through comparative phenotypic analyses (Rétaux et al., 2016). Our results bring new insights for our understanding of what is the eyefield in terms of size, shape, molecular identities and limits. The eyefield and the prospective telencephalon show unanticipated degrees of regionalization, with different subdomains of distinct cell identities and peculiar zones at the frontiers with prospective telencephalon and hypothalamus, which show significant variations in the natural cavefish mutant.

## Results

### Viewing eyefield shape and size in 3D with *rx3*

*Rx3* is a specific marker of the eyefield in the anterior neural plate (ANP) and a determinant of the retinal fate (Stigloher et al., 2006). To date, most reports of eyefield size and shape have used 2D imaging (Figure 1A-D). In *Astyanax, Rx3* expression delineated a bean-shaped domain, convex anteriorly and concave posteriorly, of smaller area in cavefish compared to the wild type surface fish embryos (Figure 1A-D; 1K-L). 3D rendering revealed distinct ventral contours in the two morphs with a midline notch partially separating the eyefield in two lobes posteriorly, which did not appear in 2D analyses (Figure 1E-H and Supplementary Figure 1). The *rx3* domain showed a 25% reduction of volume in cavefish (Figure 1M), which may correspond to the lack of posterior medial parts (Figure 1G, H, J-J’’’).

**Figure 1:**
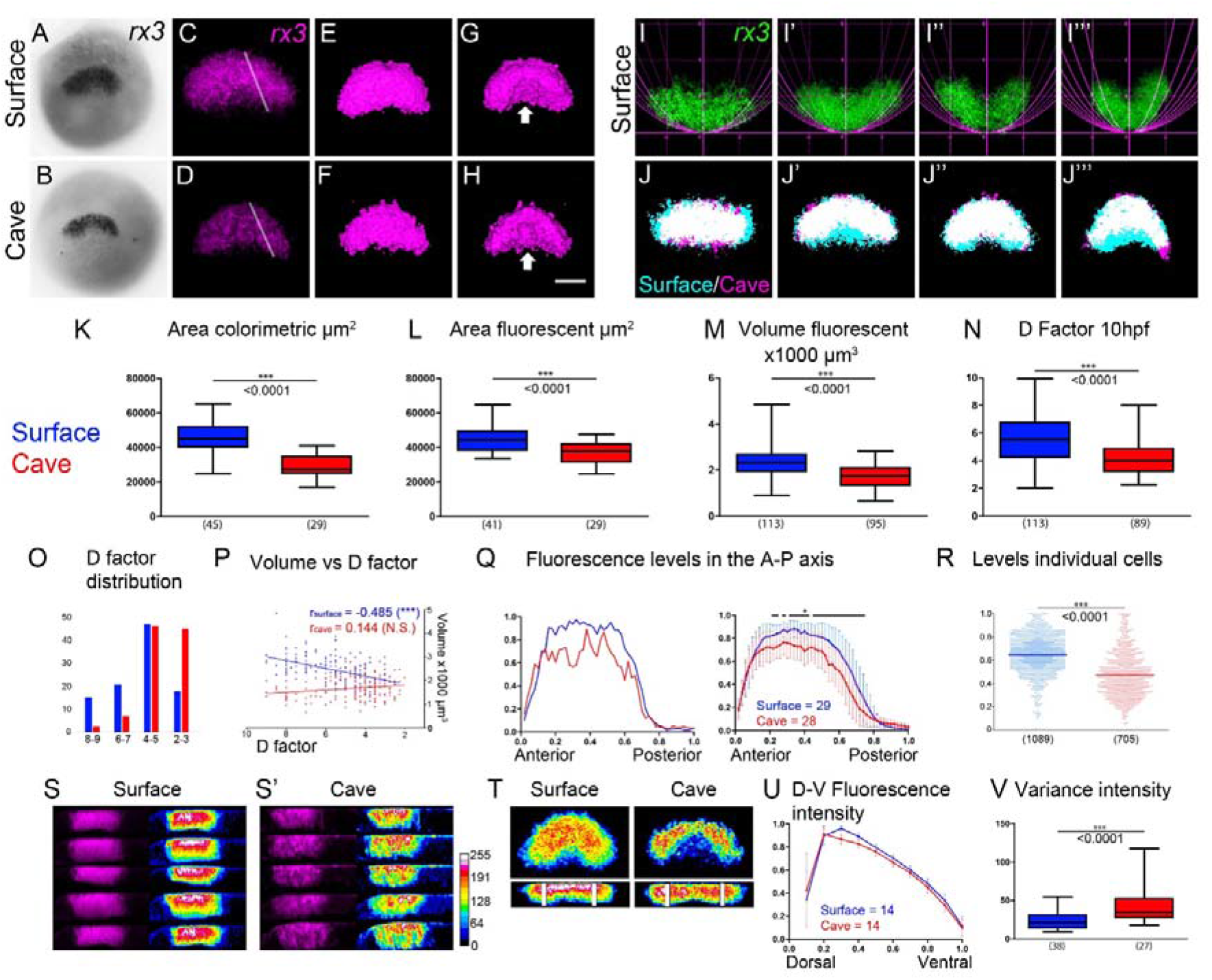
Characterization of *rx3* eyefield expression pattern and neural staging method. (A–B) *rx3* expression in surface fish and cavefish shown by colorimetric ISH. (C-H) Confocal projections showing *rx3* expression in surface fish (C, E, G) and cavefish (D, F, H). (C-D) Full stack maximum projections. (E-H) Dorsal and ventral (E, F and G, H, respectively) views from 3D reconstructions. Arrows indicate the ventral medial *rx3* depletion. (I-I’’’) Confocal full stack maximum projections showing *rx3* expression domain (green, anterior downward) in surface fish embryos combined with parabolas of different vertical scaling factors (a=1/d, d factor = 1 to 8). (J-J’’’) Confocal full stack maximum projections showing merged surface fish (cyan) and cavefish (magenta) *rx3* expression domain on specimens of similar d factor. (K,L) Surface area delineated by *rx3*-expressing domain in colorimetric ISH and after full stack maximum projections. (M) Volume of *rx3*-expressing domain. (N) Mean d factor for embryos fixed at theoretical 10hpf. (O) D factor distribution from low (8-9) to highly (2-3) curved ANP, in the two morphs. (P) Volume of *rx3* domain according to d factor. (Q) Line histogram quantification of relative pixel intensity in two representative samples (left) and averaged in many embryos (right), from lines in C,D. (R) *rx3* levels in individual cells. (S, S’) Confocal reconstructions (3 µm, from line in C and D) showing the expression of *rx3* in magenta and 16-colours ramp in surface fish (S) and cavefish (S’). (T) Confocal dorsal (half stack, top) and transversal full resliced stack (bottom) projections showing the concentric expression of *rx3* in pseudo-colours in surface fish (left) and cavefish (right). (U) Quantification of *rx3* relative levels along the D/V axis (see rectangles in T, bottom). (V) Quantification of *rx3* relative intensity variance (rectangles in T). All embryos are at 10 hpf, anterior upwards unless otherwise indicated (I-I’’’). All pictures are from flat mounted embryos, except A and B, which are whole-mounted embryos. Mann-Whitney tests were done in K-N, R and V, p values are indicated for each case. Spearman correlation were performed in P, Spearman r values are indicated. Two-ways ANOVA test was done in Q and U, black bars indicate points with significant differences (*).

### Eyefield shape and size changes are highly dynamic

Importantly, we noticed a significant dispersion of eyefield shape and volume between samples, all fixed upon their developmental time and global tailbud morphology at “theoretical” 10hpf (Figure 1I-I’’’, J-J’’’, Movie 1). To follow eyefield size and form according to time, we decided to stage ANP *a posteriori* by estimating the state of advancement of neurulation for each sample. Since the anterior curvature of ANP resembles a parabola, we designed a simple method to assign each 10hpf sample with a coefficient (d), inverse of the vertical scaling factor for a parabola. This allowed classifying samples in various subgroups according to their intrinsic morphogenesis advancement (Figure 1I-I’’’). Using this method, we found a significant shift towards higher curvatures (lower d factor) for cavefish embryo distribution (Figure 1N,O). This suggested a heterochrony in neurulation progression or onset between the two morphs, with cavefish embryos being slightly more advanced than surface fish. To compensate for this bias in sample distribution at 10hpf, we included older, 10.5hpf surface fish samples in the dataset. This allowed us to compare eyefield size within the same ranges of d factor for the two morphs (d2-3 to d8-9). Whatever the range of d factor considered, the eyefield volume was always smaller in cavefish, confirming the trend observed at 10hpf with pooled embryos (Figure 1P). In addition, we found a striking difference for *rx3*+ tissue morphogenesis between the two morphs when comparing size according to d factor. While a progressive 30% reduction of *rx3*+ eyefield volume was observed for surface fish along the progression of neurulation (Figure 1P), the *rx3*+ volume was constant within the same interval for cavefish (R^2^ = 0.02). The size of the *rx3*-expressing ANP region was thus smaller in cavefish from the start and did not condense like in surface fish embryos at the onset of neurulation.

### Expression of *rx3* within the eyefield and its defects in cavefish

Previous analyses showed that *rx3* transcript levels at 10hpf are lower in cavefish than in surface fish (McGaugh et al., 2014), which we confirmed by RNAseq and fluorescent *in situ* hybridization (data not shown and Figure 1Q). Here, we further investigated the intrinsic pattern of *rx3* expression. To achieve comparisons of spatial distributions of pixel intensities within samples, we normalized intensities from image stack acquisitions to reveal optimized pixels intensities (see materials and methods). Strikingly, *rx3* signals distributed heterogeneously in the eyefield of cavefish embryos as compared to the homogeneous distribution in surface fish (Figure 1Q). Accordingly, cavefish eyefield cells showed a wider range of *rx3* fluorescent signals than surface fish, as shown by higher dispersions of pixel intensities (Figure 1R) and variance (Figure 1V). This fully penetrant, Swiss cheese like phenotype (n=150) was prominent in the whole structure along the three axes (Figure 1C-D, 1S,S’,T). Thus, in contrast to surface fish, distinct patches of cells, even adjacent, expressed highly variable levels of *rx3* (low to high) in cavefish.

At the eyefield scale, *rx3* signals in surface fish ANP also defined concentric asymmetric gradients both in the A/P and D/V axes as revealed by false colour coded images (Figure 1S, S’ and T). We observed this pattern at every confocal plane, hence it was not due to maximum intensity projection artefact (Supplementary Figure 1). We also observed this pattern in cavefish to a lesser extent.

In summary, the *rx3* expression domain appeared more patterned and complex than previously thought and showed variations in size, shape, expression intensity and homogeneity aspect between the two *Astyanax* morphs.

### *Cxcr4b* identifies a “core” anterior subdomain within the *rx3* positive eyefield

In zebrafish, *cxcr4a* functions in the segregation of optic vesicles and telencephalon during ANP morphogenesis, and its expression depends on *rx3* (Bielen and Houart, 2012). Since *rx3* expression was strongly affected in cavefish, we reasoned that this chemokine receptor-encoding gene might be abnormally expressed in cavefish, potentially explaining optic morphogenetic defects. We cloned the *cxcr4a* and *cxcr4b Astyanax* cDNA orthologues (Supplementary Figure 2) from our EST library and analysed the *crxcr4b* probe only, since no *cxcr4a* transcript were detected at tailbud. At this stage in *Astyanax*, the *cxcr4b* expression domain was bean-shaped, was included within the *rx3*+ eyefield in both morphotypes (Figure 2A-D, Movie 2), and was 25% smaller in cavefish (Figure 2H). The *cxcr4b* domain mapped to the anterior dorsal *rx3*+ region (Figure 2A-F), identifying a core domain of co-expression, whose size was about half that of the *rx3* volume in both morphs. This core region was surrounded laterally, posteriorly and medio-ventrally by cells expressing only *rx3* (Figure 2A-F; Movie 2), thus highlighting further concentric subdivisions of the eyefield (Figure 1S,S’,T).

**Figure 2:**
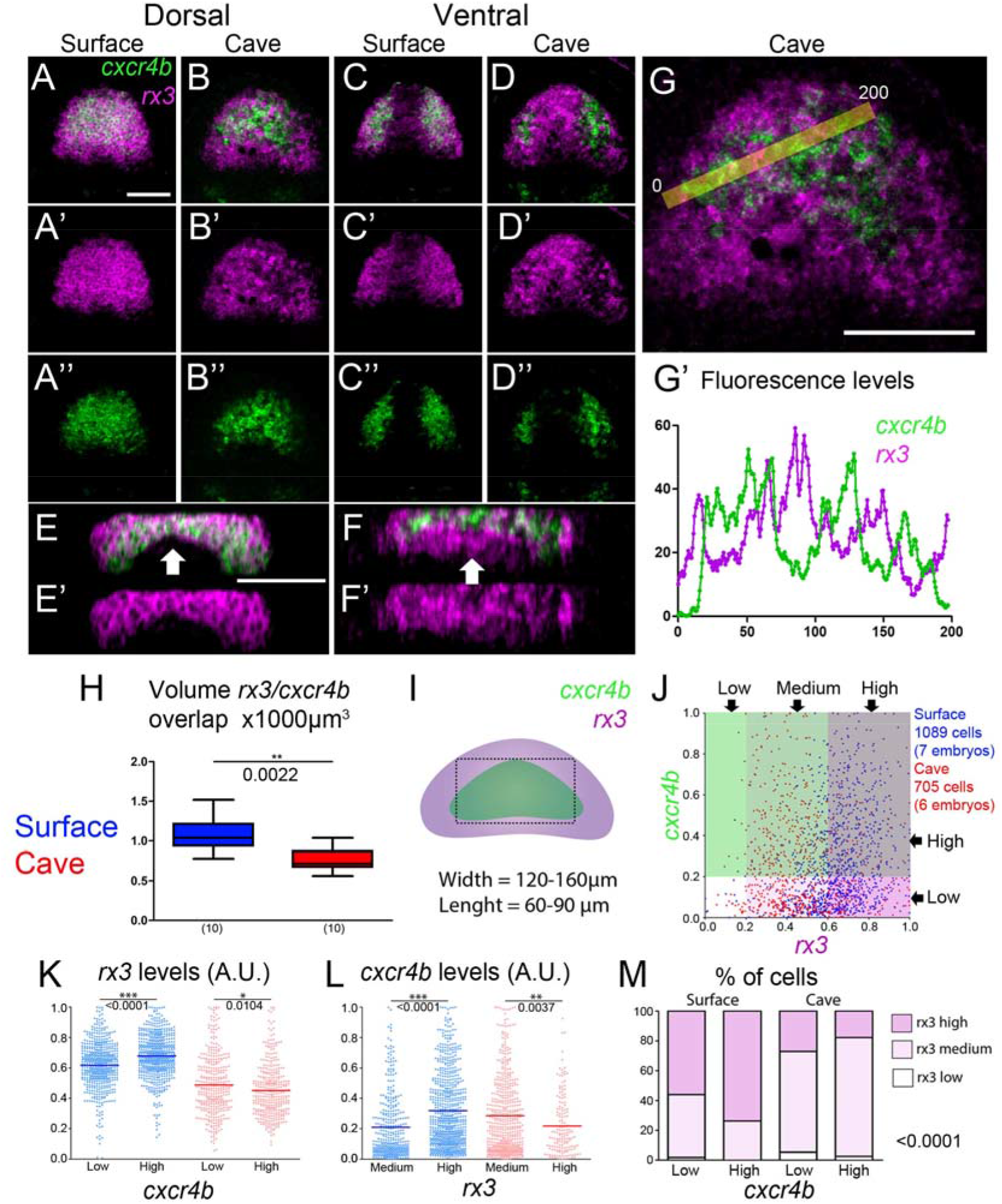
Characterization of *cxcr4b* expression in relation with *rx3* in the cavefish mutant. (A–D) Dorsal and ventral confocal maximum projections showing the expression of *cxcr4b* (green) and *rx3* (magenta) in surface fish and cavefish. (E-F) Maximum resliced stack projections in anterior frontal views of surface (left) and cavefish (right). The arrows point to a larger ventral midline domain expressing only *rx3*, in cavefish. (G) Higher magnification of embryo shown in B. (G’) Line histogram quantification of relative pixel intensity levels for the two markers (yellow bar in G). (H) Volume of *rx3*/*cxcr4b* co-expressing domain. (I) Scheme depicting *cxcr4b* and *rx3* domains relative positions and the area (dotted line) used for “single cell” quantifications shown in J-M. (J) Distribution of cells according to *rx3* and *cxcr4b* relative levels. The different areas are colour coded in relation to the expression threshold for each marker (0.2 A.U.). (K-L) *rx3* relative pixel intensity levels, according to low (<0,2 AU) or high (>0,2 AU) *cxcr4b* levels (K); and *cxcr4b* relative pixel intensity levels, according to medium (>0,2 and <0,6 AU) or high (>0,6 AU) *rx3* levels (L) in surface fish (blue) and cavefish (red). Mean values are indicated by bars. (M) Distribution of cells expressing low, medium, or high *rx3* levels according to *cxcr4b*, below or above threshold (0,2 AU). Mann-Whitney test were done in H, K and L, p values are indicated. Chi-squared test was performed in M, p value is indicated.

### *Rx3* low-expressing cells express higher levels of *cxcr4b* in cavefish

As mentioned above, *rx3* expression pattern showed drastic alteration in cavefish and lacked homogeneity, as shown by high dispersion of pixel intensities (Figure 1). It became evident that the expression patterns/levels of *cxcr4b* and *rx3* were complementary in the core region in cavefish (Figure 2B-B’’, F-F’’). Line histograms quantifications confirmed that *rx3*-low zones were *cxcr4b*-*high* and *vice versa* (Figure 2G-G’). We next quantified pixel intensities in individual cells (Supplementary Figure 3) and plotted relative levels of *rx3* and *cxcr4b* observed in each cell (Figure 2I,J). In surface fish, cells exhibiting high *cxcr4b* levels (threshold > 0.2) displayed significantly higher *rx3* levels as compared to cells with very low *cxcr4b* signals (<0.2) (Figure 2K, blue). The opposite was observed within cavefish cells (Figure 2K, red), with lower *rx3* levels in cells with high *cxcr4b* expression. We also compared *cxcr4b* levels in cells with high or medium *rx3* expression (Figure 2J). In surface fish, cells with higher *rx3* signals show higher levels of *cxcr4b* (Figure 2L, blue), whereas in cavefish cells with higher rx3 show lower levels of *cxcr4b* (Figure 2L, red). At such population scale, distributions were markedly different in the two morphs, highlighting potential opposite regulatory interactions between rx3 and *cxcr4b* (Figure 2M).

### Shape and size of the *pax6* domain

We next used the prominent eye-gene marker *pax6*, which forms together with *rx3* and *six3* the eyefield specific transcription factor network (Sinn and Wittbrodt, 2013). At the tailbud stage, *pax6* delineated bilateral triangular-shaped domains that were anteriorly and dorsally connected at the midline and showed no expression at the posterior midline, hence markedly different from *rx3* (Figure 3A, A’, B, B’). In addition to previous descriptions, dorsal and ventral projections highlighted *pax6* domain shape in 3D (Supplementary Figure 4, Movie 3, (Macdonald et al., 1997; Nornes et al., 1998; Strickler et al., 2001)). The volume of the *pax6* domain was similar in the two morphs (Figure 3I). *Pax6* partially overlapped with *rx3* anteriorly and distributed posterior to it (Figure 3C,D). We estimated the size of the *pax6+/rx3+* overlapping region and found no difference between the two morphotypes (Figure 3L), and little difference for the solo *pax6*-expressing posterior region which was slightly smaller in cavefish (not shown).

**Figure 3:**
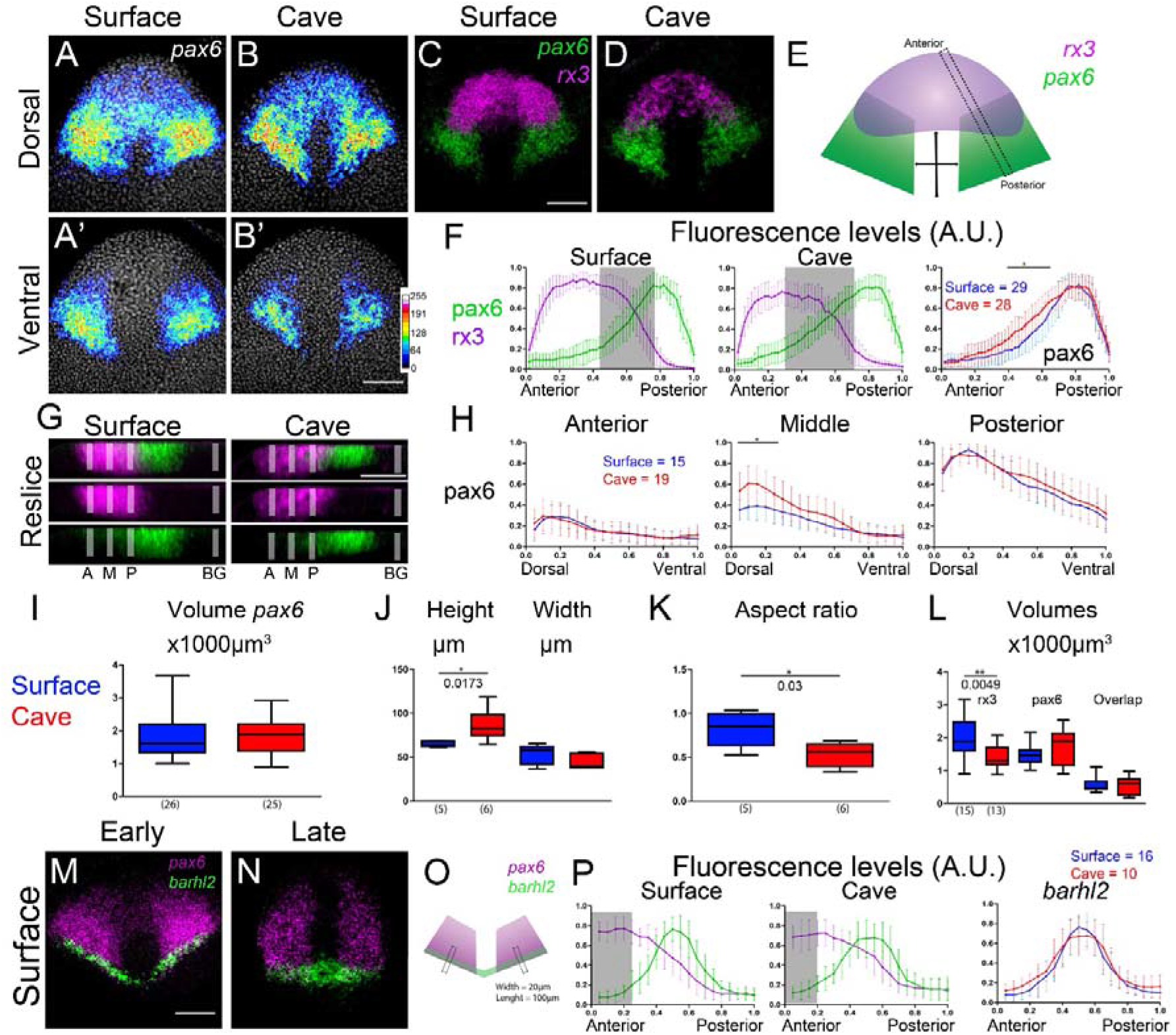
Characterization of *pax6* eyefield expression pattern and relationships with *rx3*. (A–B’) Confocal projections showing dorsal and ventral *pax6* expression in surface fish and cavefish. Color scale bar in (B’) indicates relative pixel intensities. (C-D) Full stack maximum projections showing posterior *pax6* (green) and anterior *rx3* (magenta) in surface fish (C) and cavefish (D). (E) Schematics of line histogram quantification for F and G (dashed line from anterior to posterior) and measurement of medial distances for I and J (double arrows). (F) Line histogram quantifications of relative pixel intensity levels for *pax6* and *rx3* in the anteroposterior axis. The shaded rectangle indicates fluorescence levels above threshold for both markers (>0.2 A.U.). (G) Confocal re-slice projections showing *pax6* (green) and *rx3* (magenta) expression along the D/V axis. Vertical bars correspond to anterior (A), mid (M) and posterior (P) positions and background (BG) for D/V quantifications within (A, M, P) and outside (BG) the *rx3* domain. (H) Line histogram quantifications of relative pixel intensity levels for *pax6* according to A, M, P positions (G). (I) Volume of *pax6*-expressing domain. (J) Quantifications of height and width of the *rx3*/*pax6* non-expressing domain size (according to E). (K) Quantification of height/width aspect ratio. (L) Volume of *pax6*/*rx3* co-expressing domain. (M-N) Confocal projections showing mid-dorso/ventral *pax6* and *barhl2* expression in surface fish at two different neural stage at 10hpf. (O) Schematics of line histogram quantification. (P) Line histogram quantifications of relative pixel intensity levels for *pax6* and *barhl2* (according to Q). The shaded rectangle indicates the posterior limit of the eyefield. All embryos are at 10hpf, anterior upwards. All pictures are from flat mounted dissected ANP, anterior to the top. Mann-Whitney tests were done in I-L, p values are shown. Two-ways ANOVA test were done in F, H and P, black bars indicate points with significant differences (*). Scale bars: 100 µm.

In the medial ANP, the region devoid of both *pax6* and *rx3* expression (Figure 3E) was 25% longer in cavefish (Figure 3J; d factor 3.5-4.5) but its width was similar on the same samples (Figure 3J). The resultant 1.6 fold increased aspect ratio (AR= height/width) of this *pax6*-/*rx3*-midline region in cavefish (Figure 3K), together with the differential ventral midline shape (Figure 3, Movie 3), strongly suggested that this ventral medial ANP domain was larger in cavefish. Altogether, these results strengthened the hypothesis that the reduction of cavefish *rx3* domain size concerns its medial and posterior part.

### *Pax6* expression shows anteroposterior and dorso-ventral regionalization gradients

*Pax6* transcripts distributed in a gradient manner along the anteroposterior axis, with posterior cells showing higher levels than anterior ones (Figure 3A,B,F). *Pax6* fluorescence intensity in posterior *rx3*+ cells was six times higher than in anterior *rx3*+ cells, similarly in both morphotypes (Figure 3F). Also, *pax6* levels in the intermediate region were significantly higher in the cavefish eyefield, suggesting, again, variations in the fine control of expression of this factor between the two morphotypes (Figure 3F, right).

*Pax6* expression in the ANP also showed polarization along the D/V axis (Figure 3G,H). Resliced lateral projections (rectangle in Figure 3E) were used to quantify *pax6* pixel intensities at four different positions along the A/P axis using line histograms. We drew three lines within the *rx3*-expressing domain (Anterior, Middle and Posterior; A, M, P, respectively) and a fourth outside the *pax6* domain (BG), which allowed calculating the background noise for each sample (Figure 3G) (*see materials and methods). Similar curves depicted similar D/V gradients in the anterior, middle and posterior regions of the *pax6+ rx3+* domain (Figure 3H). In each of the three regions, dorsal eyefield cells expressed higher levels of *pax6* than ventral cells. In middle sections, higher *pax6* levels were observed dorsally in cavefish embryos (Figure 3H, middle), similarly to differences observed in anteroposterior measurements (Figure 3G, right).

Altogether, these data uncovered several additional levels of eyefield regionalization. The asymmetric and gradient expression profiles of *pax6* along the A/P and D/V axes, together with the concentric gradient of *rx3* pattern, highlight unsuspected levels of combinatorial specification and distinct cell identities within the ANP, which are subtly affected in cavefish.

### Defining the posterior limit of the eyefield with *barhl2*

Since both eyefield markers *rx3* and *pax6* overlapped only partially, we asked whether the limit of the eyefield extended posteriorly to *rx3*. Staudt and Houart (Staudt and Houart, 2007) showed that the diencephalic marker *barhl2* is located directly posterior to the eyefield in zebrafish tailbud embryos. We used *barhl2* together with *pax6* to define the posterior eyefield border. Similar to zebrafish, *barhl2* co-labelled several rows (up to 8) of *pax6*+ cells at the posterior border of the *pax6* domain at different stages of neurulation, similarly in both morphotypes (Figure 3M-P). Positioning the anterior limit of the presumptive diencephalon with respect to *pax6* allowed deducing that at least two large domains subdivided the eyefield along the anteroposterior axis: an anterior domain containing cells expressing *rx3* with graded levels of *pax6*, and a posterior domain populated of cells expressing only *pax6* at high levels. These results confirmed that the eyefield is not a single territory but a composite tissue.

### The ventral limit of the eyefield: relative size and position of the hypothalamus

At the tailbud stage, the prospective hypothalamus lies just above the prechordal plate and beneath the eyefield (England et al., 2006; Pottin et al., 2011; Varga et al., 1999; Wilson and Houart, 2004; Yamamoto et al., 2004). To better characterize the ventral medial frontier of the eyefield, we thus used the hypothalamus marker *nkx2.1a* (Menuet et al., 2007; Yamamoto et al., 2004). 2D analysis using colorimetric *in situ* hybridization confirmed expansion of the *nkx2.1a* domain in cavefish (not shown; (Menuet et al., 2007; Yamamoto et al., 2004)). We next estimated the volume of the prospective *nkx2*.1a+ hypothalamus and found a significant increase in cavefish (Figure 4E).

**Figure 4:**
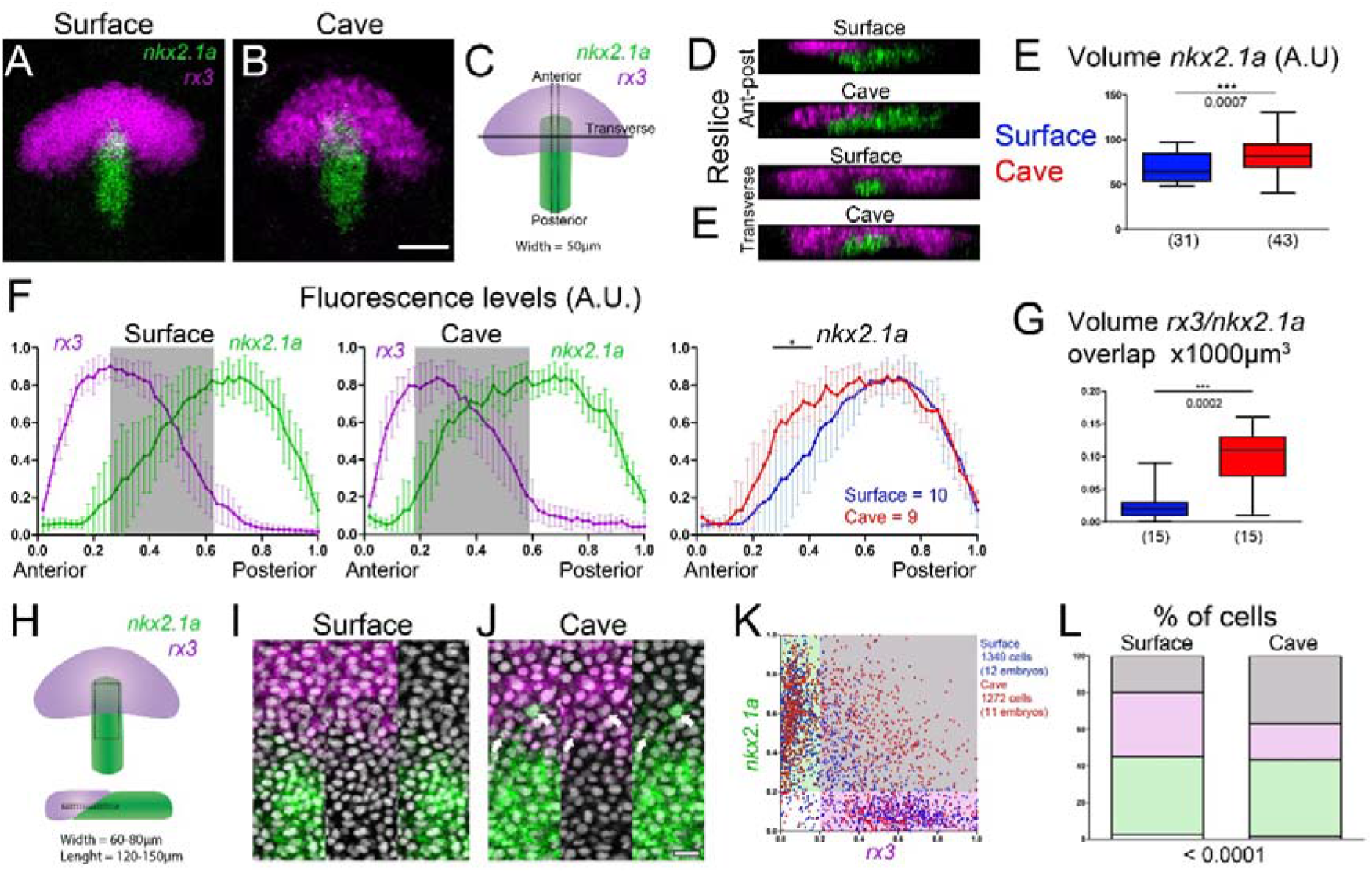
Characterization of *nkx2*.1 hypothalamic expression pattern and relationships with *rx3*. (A-B) Full stack maximum projections showing medial *nkx2.1a* (green) and *rx3* (magenta) in surface fish and cavefish. (C) Schematic of line A/P histogram quantification in F and reslices shown in D and D’. (D) Confocal re-slice longitudinal and transverse projections showing *nkx2.1* (green) and *rx3* (magenta) expression along the D/V axis in surface fish and cavefish. (E) Volume of *nkx2.1*-expressing domain. (F) Line histogram quantifications of relative pixel intensity levels in the A/P axis for *nkx2.1* and *rx3* according to C (rectangle). The shaded rectangle indicates fluorescence levels above threshold for both markers (>0.2 A.U.). (G) Volume of *nkx2*.1/*rx3* co-expressing domain. (H) Schematics of area (top) and D/V positioning (bottom) of ROIs used for single cell quantifications (I-L). (I-J) Confocal 3 µm projections showing *nkx2.1* (green), *rx3* (magenta) and nuclei (grey) in the ROI (H). (K) Distribution of single cells according to pixel relative intensities for *nkx2.1a* and *rx3* (thresholds 0.2 A.U.). (L) Frequencies of cell populations shown in (K). Mann-Whitney tests were done in E and G, p values are shown. Two-ways ANOVA test was done in F, black bar indicate points with significant differences (*). Chi-squared test was performed in L, p value is indicated. Scale bars: 100 µm (B), and 20 µm (J).

We next studied the relative spatial arrangement of *rx3* and *nkx2*.1a (Figure 4AB) and found no correlation between eyefield shape (according to d factor) and the position of the prospective hypothalamus along the A/P axis (Supplemental Figure 5). Interestingly, the prospective hypothalamus at 10hpf had a more anterior position in cavefish (Figure 4C,F), showing yet another heterochrony in the cavefish neural plate morphogenesis and prompting us to analyse the *rx3*/*nkx2.1a* frontier zone.

**Figure 5:**
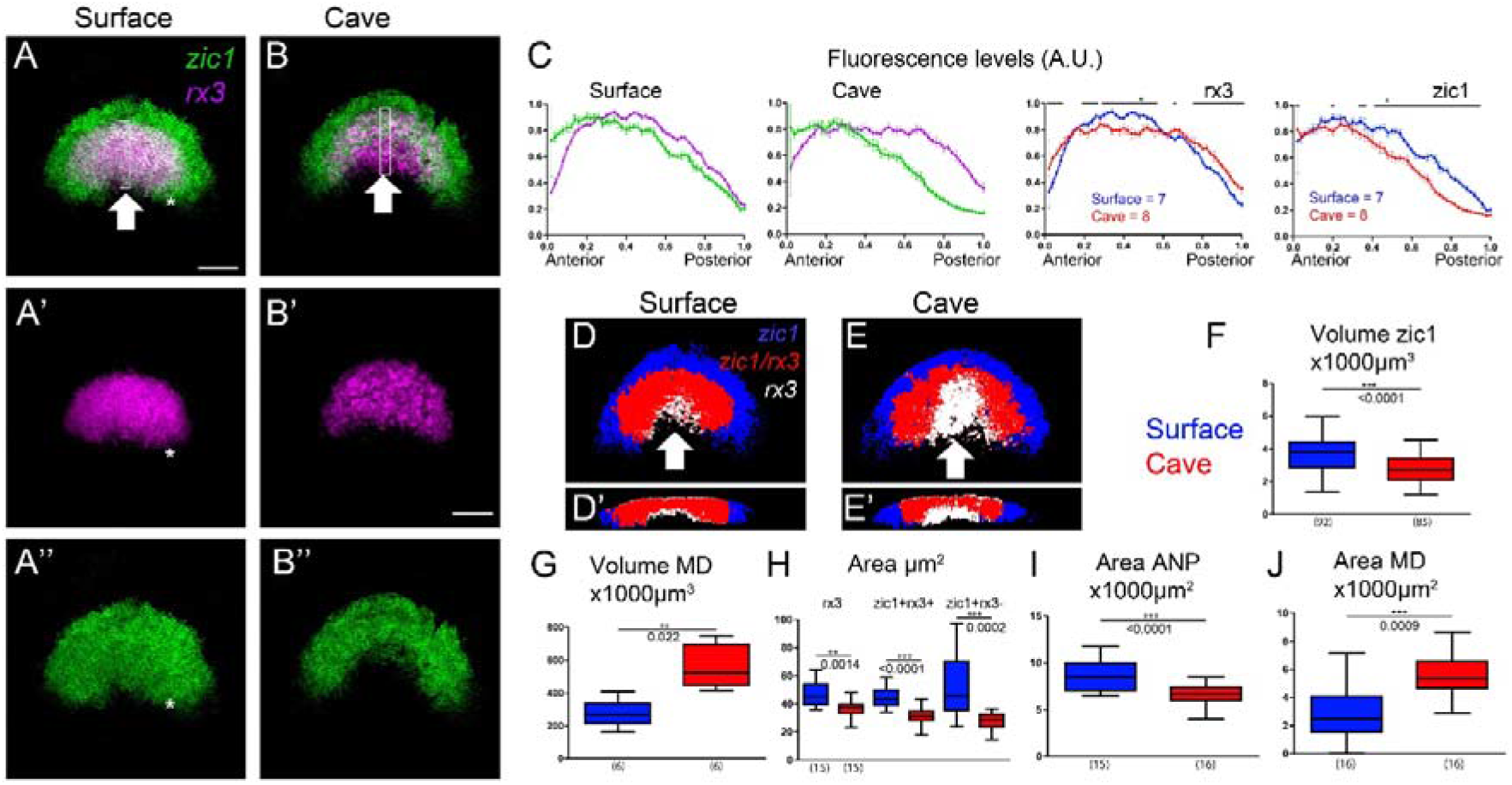
Characterization of *zic1* expression and identification of eyefield subdivisions. (A–B’’) Full stack maximum projections showing *zic1* (green) and *rx3* (magenta) in surface fish and cavefish. (C) Line histogram quantifications of relative pixel intensity levels for *zic1* and *rx3*, according to white rectangles in A, B. (D-E) Binarized images of mid-stack confocal projections showing segmented domains expressing only *zic1* (blue), *zic1* and *rx3* (red) and only *rx3* (white). (F) Volume of *zic1* expressing domain. (G) Volume of *rx3*-only expressing midline domain (MD). (H) Relative proportion of surface areas shown in D and E for total *rx3*+ eyefield domain (red and white), co-expression domain (rx3+zic1+) and telencephalic *zic1* solo-expressing domain (blue). (I) Total surface area of *zic1*-expressing + *rx3*-expressing domains as a proxy of ANP size, after mid-stack maximum projections (J) Quantification of surface area delineated by *rx3*-only expressing domain after mid-stack maximum projections (white domain in D, E). All pictures are from flat mounted dissected ANP, anterior to the top. Two-ways ANOVA test was done in C, black bars indicate zones with significant differences (*). Mann-Whitney tests were done in F-J, p values are shown.

### *Nkx2.1a* and *rx3* reveal mixed cell identities at the cavefish hypothalamus/eyefield boundary

The *rx3*/*nkx2.1a* boundary zone was studied in samples where at least one third of the *nkx2.1a* domain was advanced beneath the eyefield. In surface fish, *nkx2*.1a and *rx3* delineated two separated 3D domains, which in most cases presented a medial contact surface (n=13/15) with little or no overlap (Movie 4). A clear gap separated the *rx3* and *nkx2.1a* domains medially in few embryos (n=2/15) and laterally in all embryos (n=15/15) (Figure 4DE), suggesting an additional, unknown molecular identity in this thin tissue layer. Conversely, in cavefish the two domains were always in close contact (Figure 4DE), suggesting co-expression of *rx3* and *nkx2.1a* in the cavefish natural mutant. Indeed, a large *rx3*/*nkx2.1a* overlapping domain was found in cavefish samples (Figure 4G). “Single cell” analysis confirmed the presence of about 20 cells co-expressing the two markers per cavefish neural plate, i.e., 4 times more than in surface fish (Figure 4H-L).

At the tissue level, D/V reconstructions indicated that these cells were mainly located at the *rx3*/*nkx2.1a* boundary zone, but some were found as well at more dorsal locations within the *rx3* expressing domain, suggesting aberrant specification (Figure 4E, 4IJ). Altogether, these results revealed the existence of a very small subset of *rx3*+/*nkx2*+ cells in the surface fish neural plate at tailbud stage and their increased proportion in cavefish. Since the cavefish prechordal plate displays *Shh* expression intensification at this embryonic stage (Pottin et al., 2011; Torres-Paz et al., 2019; Yamamoto et al., 2004), the 4-fold increase in the population of *rx3*/*nkx2.1* co-expressing cells in cavefish may be a direct consequence of increased Shh ventral midline signalling.

### Shape and size of the *zic1* domain

We reasoned that *zic1*, which also responds to midline signalling in zebrafish forebrain (Maurus and Harris, 2009) and is expressed in retinal precursors at tailbud stage (Devos et al., 2019; Grinblat et al., 1998; Varga et al., 1999) would help further characterize the eyefield molecularly. At tailbud stage in *Astyanax, zic1* expression delineated an anteriorly convex bean-shaped domain, similar in shape but more indented ventro-medially than *rx3* (Figure 5AB, arrow; Movie 5). The *zic1* domain was larger than *rx3* for both morphotypes (Figure 5A-B’’; compare Figures 1M and 5F) and this was true for each d factor range considered (not shown). The *zic1* domain showed a 25% reduction of volume in cavefish when compared to surface fish (Figure 5F), which likely corresponded to the lack of posterior medial parts (Figure 5AB, arrows). We also found a 35% progressive reduction of *zic1* domain volume for surface fish along the progression of neurulation, whereas the *zic1*+ volume was constant within the same interval for cavefish (data not shown). As shown for *rx3*, the size of the *zic1*-expressing ANP region did not condense like in surface fish embryos at the onset of neurulation.

### *Zic1* identifies two major domains in the eyefield

*Zic1* expression partially overlapped with that of *rx3*, thus identifying combinatorically three zones within the ANP (Figure 5AB). Anteriorly, *zic1* extended beyond *rx3* in both morphs (Figure 5DE, blue, Movie 6), in a zone corresponding to *emx3*+ telencephalic precursors (see Figure 7). The size of this *zic1*+/*rx3*-telencephalic region was 30% smaller in cavefish whatever the d factor considered (Figure 5H). A second zone was composed of *zic1*+/*rx3*+ cells, corresponded to a sub-territory of the *rx3* domain, and was also smaller in cavefish (Figure 5DE, red; Figure 5H). The third, medial region, was populated with *rx3*+/*zic1*-cells (Figure 5DE, white and arrowheads) and was twice larger both in surface area and volume in cavefish embryos (Figure 5G,J, Movie 6). This domain was four-fold larger in size proportion relative to either *zic1* or *zic1*+/*rx3*+ domains in cavefish.

**Figure 6:**
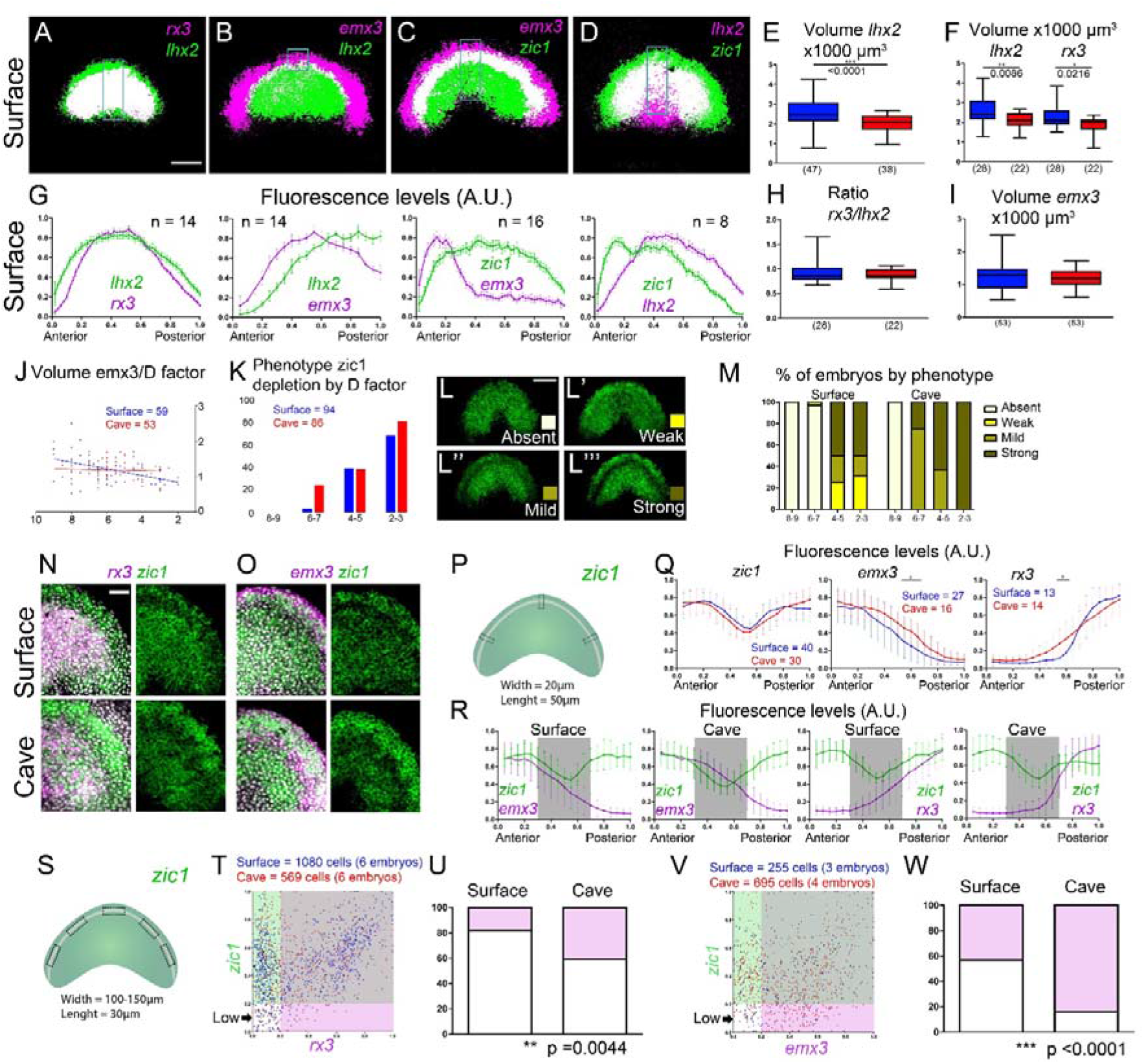
Identification of telencephalic subdivisions and characterization of the eyefield/telencephalic transition zone. (A-D) Images obtained after maximum intensity projection and binarization of individual channels showing expression of the indicated markers in surface fish at 10hpf. (E) Volume of *lhx2* expression domain. (F) Volume of *lhx2* and *rx3* used for ratio calculation in H. (G) Profiles of fluorescence intensity in the anteroposterior axis in surface fish embryos as indicated in A-D (rectangles). (H) Ratio of *rx3*/*lhx2* volumes. (I) Volume of *emx3* expression domain. (J) Volume of *emx3* domain according to d factor. (K) Frequency of the *zic1* depletion phenotype according to different d factors. (L-L’’’) Expressivity of the *zic1* depletion phenotype shown in 4 categories, absent (L), weak (L’), mild (L’’) and strong (L’’’). (M) Percentage of embryos of each *zic1* depletion pattern category (L-L’’’) for different d factors in surface fish and cavefish. (N) *rx3* and *zic1* expression in surface fish and cavefish. (O) *emx3* and *zic1* expression in surface fish and cavefish. (P) Scheme of the selection of ROIs used to generate the fluorescence plot profiles shown in Q-R. (Q) Comparison of fluorescence plot profiles at the *zic1*-depletion zone between cavefish and surface fish for *zic1, emx3* and *rx3*. (R) Fluorescence plot profiles at the *zic1*-depletion (indicated by shaded area) for *zic1, emx3* and *rx3* in surface fish and cavefish. (S) Scheme of the selection of ROIs used for the quantification of fluorescence in single cells in zones of low *zic1* expression levels. (T,V) Distribution of cells according to *rx3* and *zic1* (T) and em3 and *zic1* (V) relative levels. The 0.2 value is considered as the expression threshold for each marker, distinguishing low and high levels. (U,W) Percentage of cells with high (pink) or low (white) *rx3* (U) or *emx3* (W) levels in cells with low *zic1* levels (AU). All pictures were obtained from flat mounted dissected ANP, visualized in dorsal view, with anterior to the top. Scale bars A-D and L = 100 µm; and N = 50µm. Mann-Whitney tests were done in E, F, H and I, p values are indicated for each case. Spearman correlation were performed in J. Two-ways ANOVA tests were done in Q, black bars indicate points with significant differences (*). Fisher exact test were performed in U and W, p values are indicated in the bottom.

**Figure 7:**
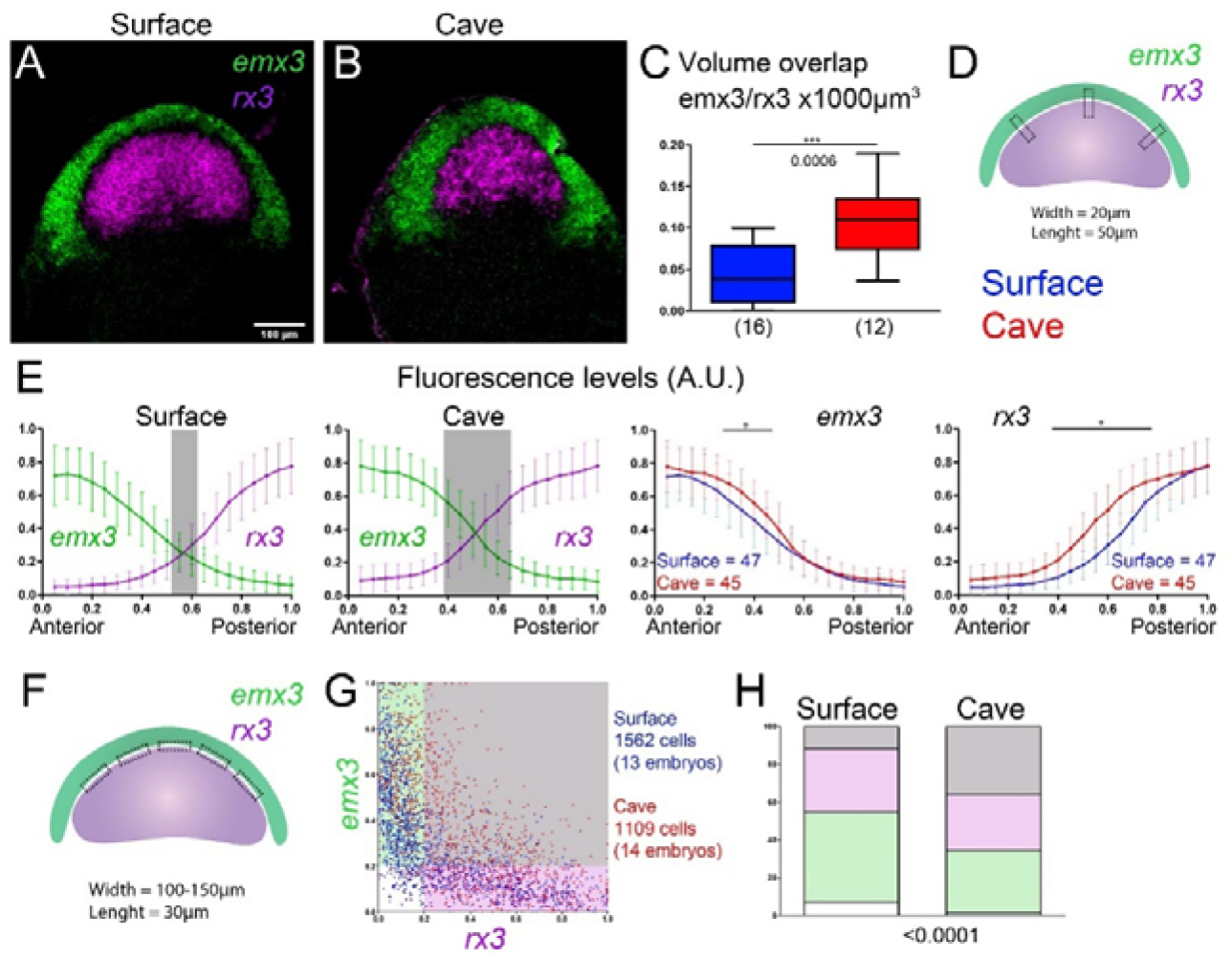
Identification of telencephalic subdivisions and characterization of the eyefield/telencephalic transition zone. (A-B) Expression of *emx3* and *rx3* in surface fish and cavefish at 10hpf. (C) Volume of overlap between *emx3* and *rx3* expression domains. (D) Scheme for the selection of ROIs used to generate the fluorescence plot profiles at the emx-*rx3* boundary. (E) Profiles of fluorescence intensity in the anteroposterior axis for *emx3* and *rx3* in surface fish and cavefish (first and second plots, respectively). The shaded rectangle indicates fluorescence levels above threshold for both markers (>0.2 A.U.). The third (*emx3*) and fourth plot (*rx3*) compare respective expression at the boundary of the two markers between surface and cavefish. (F) Scheme for the selection of ROIs used for the quantification of fluorescence in single cells at the emx-*rx3* boundary. (G) Distribution of cells according to *rx3* and *emx3* relative levels. The different areas are colour coded in relation to the expression threshold for each marker (0.2 A.U.). (H) Percentage of cells according to fluorescence levels. The same colour code is used as in G. Grey: high *rx3* and *emx3*; pink: high *rx3* and low *emx3*; green: low *rx3* and high *emx3*; white: low *rx3* and *emx3*. Mann-Whitney tests was done in C, p value is indicated. Two-ways ANOVA tests were done in E, black bars indicate points with significant differences (*). Chi square test was performed in H, p value is indicated in the bottom.

Thus, two major subdivisions arise in the eyefield based on the combination of *zic1* and *rx3* expression (Figure 5DE, red/white) and the relative proportions of these two sub-regions vary between the two morphs, suggesting a potential trade-off mechanism within the optic primordium.

### Refining eyefield lateral posterior subdivisions with *zic1*

Closer inspection of the posterior lateral eyefield revealed that *zic1* expression extended slightly posteriorly to *rx3* in the lateral region of surface fish embryos (Figure 5, asterisks; Supplemental Figure 5). 3D analysis confirmed this pattern along the entire DV axis. This additional molecular identity (e.g., *zic1*+/*rx3*-/*pax6*+ cells) indicated further compartmentalization of the eyefield lateral posterior regions. This domain was reduced in cavefish as shown by line histogram quantifications (Supplementary Figure 5), pointing to another regionalized cell specification variation in the anterior neural plate.

### *Zic1, emx3* and *lhx2* subdivide the telencephalon in three domains

Since *zic1* expression domain extended more anteriorly than *rx3*, we wished to define the anterior eyefield boundary and to characterize the presumptive telencephalon molecularly as well, suspecting subdivisions like in the eyefield. We used combinations of several markers including *zic1, lhx2* (previously reported to be differentially expressed in the cavefish ANP; (Pottin et al., 2011), *rx3* (eyefield specific) and *emx3* (telencephalon specific, (Houart et al., 1998; Morita et al., 1995)). We first used *lhx2* and *rx3* in surface fish and found that the *lhx2* anterior limit systematically exceeded that of *rx3* (Figure 6A and 6G, first plot), which was also the case for cavefish (data not shown). The *lhx2* domain showed a 20% reduction of volume in cavefish compared to surface embryos (Figure 6E), while keeping the relative volume proportions of the *rx3* and *lhx2* domains similar to surface fish (Figure 6H). Using *emx3*, whose domain size was not significantly different between the two morphs (Figure 6I), we observed a reduction in telencephalic size according to increased curvature in surface fish, as opposed to a constant volume whatever d factor considered in cavefish (Figure 6J). This trend was similar to our observations for the *rx3*+ eyefield (Figure 1P). We then confirmed that the anterior most *lhx2*-expressing cells (*rx3*-) circumferentially mapped to the posterior part of *emx3*-expressing telencephalic anlage, in the two morphs (Figure 6B and G, second plot). We also detected the *zic1* anterior limit lying posterior to the *emx3* anterior border (Figure 6C and G, third plot), in a similar way in the two morphs (data not shown). Finally, we also established that the *zic1* domain extended beyond the anterior limit of *lhx2*, similarly in the two morphs (Figure 6D and G, fourth plot). Altogether, these combinatorial mappings demonstrate that the prospective telencephalon at neural plate stage is divided at least in three subdomains, potentially corresponding to different fates (Figure 8).

**Figure 8:**
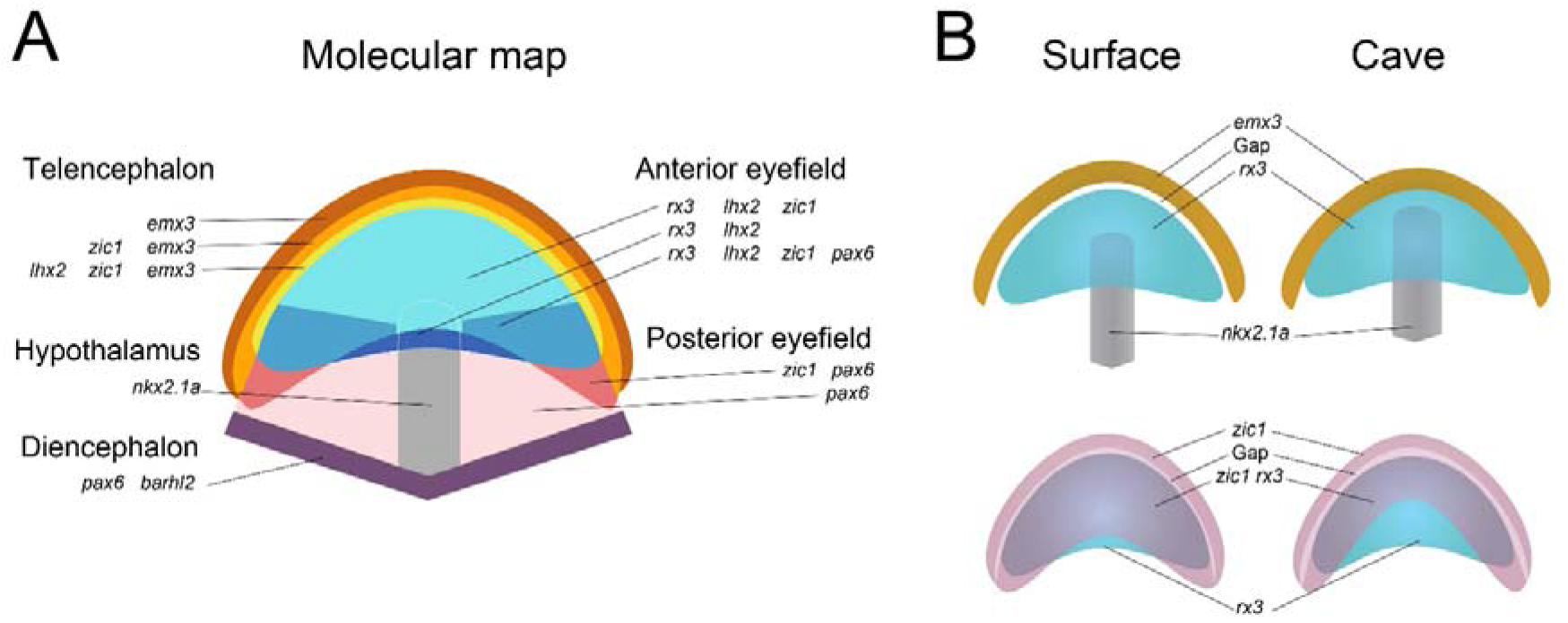
Molecular map of the ANP and modifications observed in the *Astyanax* morphs. (A) The major divisions of the ANP and further subdivisions of the eyefield and prospective telencephalon. The eyefield shows A/P regionalization. Three distinct domains of unique molecular identities constitute the anterior eyefield: *rx3+lhx2+zic1*+, *rx3+lhx2+zic1+pax6+* and *rx3+lhx2*+. The posterior eyefield shows two distinct domains: *zic1+pax6+* and *pax6+*. The posterior diencephalic limit to the eyefield corresponded to cells co-expressing *barhl2* and *pax6*. Its ventro-medial border is limited by hypothalamic *nkx2*.1a expression. Ventral anterior hypothalamus is indicated in dashed line. The prospective telencephalon divides in three distinct molecular signatures: *emx3+lhx2+zic1*+, *emx3*+ *zic1*+, and *emx3*+. (B) Comparison of salient phenotypes in the two morphotypes. (Top) A clear gap separates the eyefield and telencephalic territories in surface fish, whereas they join in the cavefish. Similarly, the ventral eyefield and dorsal prospective hypothalamus just appose in surface fish, whereas they are intermingled in cavefish. In cavefish reduction of domain separation associates with mixed identities at the boundaries. The hypothalamus in cavefish has a larger size, and it is located more anteriorly. (Bottom) The zone of *zic1* depletion (gap) at the eyefield-telencephalic boundary appears early in cavefish, and is more pronounced. The eyefield midline domain expressing only *rx3* shows expansion in cavefish.

### A “low-*zic1* crescent” in the anterior ANP, is affected in cavefish

With these anterior ANP markers at hand, we then focused on the eyefield/telencephalic boundary. Careful inspection of *zic1* pattern at 10hpf revealed the presence of a thin zone of lowered expression at the putative telencephalic/eyefield border in some samples (Figure 6L-L’’’). This phenotype was rare in surface fish (n=3/50, 6%) and more penetrant in cavefish (n=15/58, 25%) using pooled 10hpf samples of various d factors. This was reminiscent of a potential neurulation heterochrony effect in sample distribution (see Figure 1O). Accordingly, the presence of the “low-*zic1* crescent” was related to the d factor and its penetrance increased at later neurulation stages in the two morphs (d factor 4-5 and 2-3) (Figure 6K). Using a qualitative classification as absent, weak, mild or strong (Figure 6L-L’’’), we found a significant increase and full expressivity of the “low-*zic1* crescent” in cavefish, for samples with low d factor (Figure 6M). Thus, *zic1* down-regulation in this peculiar domain takes shape in a similar fashion in the two morphs but seems faster in cavefish, highlighting a differential temporal control.

To characterize molecularly the “low-*zic1* crescent”, we used combinations of markers including *rx3, emx3* and *zic1*. 3D renditions suggested that it matched the border region of the eyefield and prospective telencephalon, forming a gap between the two territories (Figure 6NO; Movie 7). Line histograms were drawn across the zone of interest in 2µm maximum projections of ANPs double stained for *zic1*/*emx3* or *zic1*/*rx3* (Figure 6P). Quantifications showed that *zic1* relative pixel intensity profiles were similar in the two morphs (Figure 6Q, left), further confirming the shared characteristics of the *zic1* depletion crescent for samples of similar morphogenetic stages. Intensity profiles of *emx3* and *rx3* showed that the “low-*zic1* crescent” had a prospective telencephalic identity in surface fish, as deduced from intermediate *emx3* levels and very low or no *rx3* expression (Figure 6R, first and third plots, respectively).

Strikingly, both the relative levels of *emx3* and *rx3* were elevated in the low-*zic1* crescent in cavefish (Figure 6R, second and fourth plots, respectively), suggesting improper cell specification in this eyefield/telencephalon intermediate region. These profiles may also be interpreted as the result of posterior and anterior shift of the *emx3* and *rx3* limits, respectively, in cavefish. Using “single cell” analysis to quantify relative pixel intensities in the *zic1*-depleted zone (Figure 6S). In this domain identified by low levels of *zic1* (indicated as “Low” in Figure 6T and 6V), a fraction of cells expressed important levels of *rx3*, corresponding to about 20% in surface and 40% in cavefish embryos (Figure 6TU). We also identified some *emx3*-expressing cells within this domain, which corresponded to ∼40% and more than 80% in surface and cavefish embryos, respectively (Figure 6V, W). This confirmed that the *emx3* posterior boundary was slightly shifted posteriorly in cavefish, in agreement with the high proportion of cells expressing *emx3* in the “low-*zic1* crescent” (80-90% in cavefish vs 40% in surface fish). Curiously, in cavefish this particular *zic1*-low region displayed higher relative levels of both *emx3* (telencephalic marker) and *rx3* (optic marker) (Figure 6QR). Altogether, these data further characterized a non-previously described zone in the ANP, showing subtle differences in cavefish, with temporal and molecular shifts of the “low-*zic1* crescent”.

### Mixed cell identities at the eyefield/telencephalon boundary in cavefish

Finally, we analysed the telencephalic/eyefield boundary proper. In surface fish, we observed a fully penetrant gap between the *rx3* and *emx3* domains (Figure 7A, Movie 8), which was independent from the neurulation stage/d factor (data not shown). Line histogram quantifications confirmed low (<0.2) relative pixel intensities at the intersection of the two curves (Figure 7D and E, first plot). Strikingly, this *emx3*-/*rx3*-gap zone never existed in cavefish (Figure 7B, Movie 8), and line histograms revealed significant higher relative pixel intensity levels for both markers at the intersection zone (Figure 7E, second plot).

Comparison of *emx3* and *rx3* curves of fluorescence intensity revealed a slight posterior and anterior shift, respectively, in cavefish compared to surface fish (Figure 7E, right plots). Accordingly, volume analyses detected a significant *emx3*/*rx3* overlapping domain in cavefish (Figure 7C). Further, “single cell” analysis at the interface zone showed a 3-fold increase in the proportion of cells expressing medium to high relative levels of both *emx3* and *rx3* (Figure 7G and H, grey) in cavefish, indicating an important degree of co-expression of the two markers. We also found in the cavefish a 5-fold decrease in the percentage of cells expressing low levels for both markers (Figure 7GH, white), consistent with the gap observed at the telencephalic/eyefield interface in surface fish but not cavefish embryos. In sum, this peculiar “gap zone” expressing lowest levels of *rx3* and *emx3* probably corresponds to another yet undescribed ANP subdivision, which again seems phenotypically affected in cavefish.

## Discussion

### What is the eyefield?

Studies in a variety of species have suggested that the vertebrate eyefield is specified by a specific combination of few transcription factors that define a single homogeneous forebrain subdivision e.g., (Varga et al., 1999; Zuber et al., 2003). Accordingly, in the literature, *pax6, lhx2* or *rx3* are considered as interchangeable markers and universal “master” eye genes, a notion which fits well with the absence of eye development and anophtalmic phenotype of knock-out/mutant animals (Gehring, 1996; Loosli et al., 2001; Porter et al., 1997). Our 3D maps reveal that the so-called eyefield is in fact already exquisitely molecularly regionalized at late gastrula stage (Figure 8). Here, using “only” five markers (*rx3, lhx2, pax6, zic1* and *cxcr4*), we could identify six subdomains, on top of which expression gradients superimpose an additional degree of patterning complexity.

Three major eyefield subdivisions in the A/P axis are given by the *rx3*/*pax6* couple: an anterior *rx3*-only, an intermediate *rx3*-*pax6*, and a posterior *pax6*-only zone. In the medio-lateral axis, the *rx3*/*zic1* couple seems to be determinant, defining a medial *rx3*-only, an intermediate *rx3*-*zic1* and small lateral *zic1*-only zone. In the dorso-ventral dimension, positional information is given by the *pax6* expression gradient, the *rx3*/*zic1* combination in the medial region and the anterior *cxcr4*-expressing core. These molecularly-defined eyefield subdomains follow the 3 embryonic axes and can integrate in a theoretical framework of forebrain development which results from potential causal signalling effects (Rubenstein et al., 1998).

The eyefield should thus be modeled as a highly regionalized structure, well before its bilateral evagination into two optic vesicles. Since eyefield cells allocate to specific positions during neurulation and formation of optic vesicles (Devos et al., 2019), and since they conserve their relative positions during eye morphogenesis (Kwan et al., 2012), it is likely that many more distinct eyefield cell identities, defined by combinatorial expression of additional transcription factors, are already specified at late gastrula.

Neuroanatomical studies have proposed the Optic Recess Region (ORR) as a morphogenetic entity organized around the optic recess, between the hypothalamus and telencephalon, at 24hpf in zebrafish (Affaticati et al., 2015). At tailbud stage, the so-called eyefield is thus fated to generate both the retinas and the derivatives of the ORR in the medial neural tube (the preoptic region). Under this framework, and thanks to the comparative analysis of surface and cavefish morphs, we can venture proposing tentative fates for some of the eyefield subdivisions we have observed. The *rx3*+/*zic1*+/*pax6*+ expressing cells potentially represent the anterior eyefield fated to become retinas, which are smaller in cavefish (Alunni et al., 2007; Devos et al., 2019; Yamamoto and Jeffery, 2000). By contrast, the medial *rx3*+/*zic1*-expressing cells, which occupy a larger territory in cavefish, could prefigure the medial ORR and the optic stalk region, which are larger in cavefish at 24hpf (Torres-Paz et al., 2019). Thus, the relative proportions of these two sub-regions identified within the eyefield would vary between the two morphs, illustrating a potential trade-off mechanism within the optic primordium, probably because of variations in signalling (Torres-Paz et al., 2019), and in agreement with previous comparative fate maps in the two morphs (Pottin et al., 2011). Such mechanism may also underlie the natural variations of fine forebrain and ocular anatomy encountered in different species of vertebrates.

Our study also highlights the subdomains of the prospective telencephalon, which, like the prospective diencephalon (Staudt and Houart, 2007) and eyefield (this study), is composed of several concentric territories of specific cell identities, most likely prefiguring distinct cell fates. Our proposed molecular map of the anterior neural plate, which integrates concentric gene expression, as previously suggested (Zuber et al., 2003) (Figure 8), will be relevant to spatially interpret future single cell transcriptomics analyses of developmental trajectories.

### Variations at the borders

#### Prospective telencephalon-eyefield boundary

*Rx3*, whose expression starts as off 70% of epiboly in zebrafish and *Astyanax* (data not shown), gradually sharpens its anterior border and is necessary for the segregation of eyefield *versus* telencephalic fates (Stigloher et al., 2006). Surprisingly, our 3D confocal analysis revealed a small, previously unreported, 2-3 cells wide circumferential territory separating prospective telencephalon (*emx3*+) and eyefield (*rx3*+) and expressing both markers at low relative levels at tailbud stage, independently of the curvature of the neural plate. In cavefish, we observed a population of cells showing a mixed identity, i.e. higher relative levels for both markers compared to surface fish. Since higher *emx3* relative levels might repress *rx3*, which is expressed at lower levels in cavefish, one reasonable interpretation would be that progenitors at the border will adopt a telencephalic fate later on. Our observation of a similar prospective telencephalon size in both morphotypes invalidates a slight posterior telencephalon expansion hypothesis, but not a potential progressive telencephalic fate adoption in cavefish. Of note, this very region also corresponds to the “low-*zic1 crescent*” we have discovered, and which is larger and has different expression dynamics in cavefish. Together, these findings suggest two, non-mutually exclusive models: i) progenitors at the border region maintain dual cell fate potentiality at tailbud, that is solved-out later-on, ii) this region constitutes an uncovered anterior subdivision of the neural plate with specific (unknown) cell fate. Signaling candidates that might be responsible for the variation observed at the telencephalic/eyefield boundary between the two morphs include Fgf and Hedgehog (Cavodeassi and Houart, 2011), which are both subtly modified in cavefish (Pottin et al., 2011; Yamamoto et al., 2004).

### Mixed identities at the hypothalamus eyefield boundary

In surface fish, the *rx3* and *nkx2*.1 expression domains are clearly non-overlapping, although several cells at the contact zone can express the double identity. Whether this condition is transient and viable is unknown, but an attractive hypothesis would be that these cells prefigure anterior/tuberal hypothalamic neuropeptidergic fates (Muthu et al., 2016). Interestingly, in cavefish the intersection between ventral midline eyefield and prospective hypothalamus territories was significant, and the cell population expressing both *rx3* and *nkx2*.1 was doubled as compared to surface fish – in line with reported variations in neuropeptidergic neuroanatomy in the basal forebrain of the two *Astyanax* morphs (Alié et al., 2018). Such increased mixed identity at this specific location in cavefish might result from signals emanating from the ventral midline. Shh, whose expression is expanded in cavefish prechordal plate at this stage (Pottin et al., 2011; Ren et al., 2018; Torres-Paz et al., 2019; Yamamoto et al., 2004) and whose manipulation affects *nkx2*.1 expression in vertebrates including cavefish (Pabst et al., 2000; Rétaux et al., 2008), constitutes a good candidate for this novel phenotype. Since Shh signalling during gastrulation controls the future numbers of specific peptidergic neuronal populations (Allié et al, 2018), it is tempting to hypothesize that these neuronal subsets might be already primed at tailbud. Lineage analyses should confirm these propositions.

### *Rx3*, an exemplary read-out of eyefield patterning and cell identity variations in cavefish

More striking than a systematic reduction of mRNA levels in cavefish as described via transcriptomics (McGaugh et al., 2014; unpublished results from our group), *rx3* expression was affected in many additional ways. Its domain of expression was smaller, potentially explaining the small size of early evaginating cavefish optic vesicles (Devos et al., 2019). Its fine regulation in co-expression with other transcription factors was modified at the boundaries of the eyefield (see above), with possible consequences on cell identities at the telencephalic and hypothalamic borders. And, last but not least, *rx3* expression level was heterogeneous at cell population level within the eyefield, showing full penetrance for this “salt and pepper” phenotype - which was not observed for the other transcription factors studied, *pax6, zic1*, or *lhx2*. Alone and in combination with other markers, *rx3* heterogeneity suggest that cells in the cavefish prospective retinas not only show reduced *rx3* levels compared to surface fish, but are also variable and unpredictable, thereby most likely affecting cell programs and behaviors differently between neighboring cells in a given sample, as well as between different samples. Finally, this cell-level *rx3* expression heterogeneity indeed seems to affect the control of the expression of effector molecules: as shown in combination with the chemokine *cxcr4b*, cells with medium *rx3* levels display high *cxcr4b* levels and are increased in proportions in cavefish. These quantifications of *cxcr4b* and *rx3* levels in the natural cavefish mutant background suggest more complex epistasis between *rx3* and *cxcr4* than previously described, and/or additional regulatory modes. In zebrafish, rx3-dependent *cxcr4a* expression demarcates the border of the eyefield with the telencephalon and confers specific segregative properties to the eyefield, which is required for correct neurulation, as soon as telencephalic cells start to move on top of the eye precursors during the early stage of keel formation (Bielen and Houart, 2012). In cavefish as well, rx3-dependent *cxcr4b* regulation might affect adhesive or migratory properties of optic cells during early eye evagination, as we have recently observed through live imaging (Devos et al., 2021).

It will be of great interest to link *rx3* defective genetic programs with eye morphogenesis and size defects in cavefish. We propose that the different aspects of *rx3* dysregulation have distinct developmental and genetic origins. The size of the *rx3* expression domain would be a “simple” consequence of the many signaling modifications that have been described in cavefish during and at the end of gastrulation (Hh, Fgf, Bmp and Wnt: Hinaux et al., 2016; Pottin et al., 2011; Ren et al., 2018; Torres-Paz et al., 2019; Yamamoto et al., 2004). The *rx3* expression levels on the other hand, and mostly its heterogeneity aspect, most probably result from intrinsic cis-regulatory changes at the level of the *rx3* locus.

In zebrafish, medaka and *Astyanax* surface fish, the *paired-type* homeodomain transcription factor *rx3* is critical for eye development, as all *rx3* mutant fish are eyeless (Kennedy et al., 2004; Loosli et al., 2003; Loosli et al., 2001; Warren et al., 2021). There, *rx3* controls survival of eye progenitors (Kennedy et al., 2004) as well as optic vesicle evagination and neuronal differentiation (Loosli et al., 2003), as revealed in the zebrafish *chokh*/*rx3* loss-of-function background. In cavefish, no major apoptosis is recorded in optic vesicle cells during eye morphogenesis (Devos et al., 2021), and retinal apoptosis starts later around 36-48hpf (Alunni et al., 2007), in a lens-dependent manner (Yamamoto and Jeffery, 2000). It is nevertheless possible that heterogeneous and low *rx3* levels could allow cavefish optic cells to survive, and prime them for future apoptotic programs. At the tissue level, *rx3* heterogeneity may participate in a potential mechanism allowing progressive and controlled degeneration.

### Heterochronic cavefish

Our 3D comparative analyses revealed substantial and more subtle differences in surface fish and cavefish neural plates, both in terms of eyefield regionalisation/subdomains and at single cell level. They also revealed striking differences in terms of apparent tissue dynamics: while some extent of eyefield condensation seems to occur at the onset of neurulation in surface fish, this global process of size reduction does not seem to take place in cavefish.

We developed a method of anterior ANP curvature normalisation to be able to compare eyefield domains shape and size for similar neural development. Importantly, and based on the “d factor”, the observed patterning differences were not subject to timing (i.e. they were observed for all d factors and were not delayed) and thus did not depend on advancement of neurulation and keel formation.

In zebrafish, during the initial step of optic vesicle evagination at the onset of neurulation, cell behaviours are complex: some eye-fated cells behave like the nearby telencephalic cells and converge toward the midline to form the neural keel, while others lag behind and keep the eyefield wide (Ivanovitch et al., 2013; Rembold et al., 2006; reviewed in Bazin-Lopez et al., 2015; Sinn and Wittbrodt, 2013; Wilson and Houart, 2004). This convergence towards the midline may underlie the decrease in volume that we observed in surface fish for both the telencephalic/*emx3* tissue and the optic/*rx3* tissue, as a function of the d factor. Such neural plate condensation seems to take place in a short period in surface fish (less than 30min) and the process seems to be impaired or absent in cavefish. Again, the intrinsic properties of the cavefish tissue, including its improper heterogeneous *rx3* expression and the consecutive dysregulation of *cxcr4* which normally influences the cohesion of eyefield cells (Bazin-Lopez et al., 2015), might be responsible for this phenotype.

## Conclusion

As expected, all the major subdivisions that we have combinatorically defined in the eyefield and telencephalon of *Astyanax mexicanus* were found in both morphotypes (Figure 8), and are likely present in other fish and other vertebrate species. Detailed fate maps, lineage analyses and pseudo-time single-cell transcriptomics analyses will decipher their respective identities and outcomes. The differences we have observed between the cave and surface morphs of our model species can guide us towards functionally relevant hypotheses for the genetic specification of vertebrate optic tissues. They also guide us towards the identification of the genetic basis of cavefish eye defects.

## Materials and methods

### *A. mexicanus* embryos

Our surface fish colony originates from rivers in Texas (USA) and our cavefish colony derives from the Pachón cave in the state of Tamaulipas (Mexico). Embryos were obtained by *in vitro* fertilization, after induction of the breeding colony for gamete maturation and reproduction by changes in water temperature (Elipot et al., 2014). The embryonic development of *A. mexicanus* at 24°C is similar and synchronous for both morphotypes (Hinaux et al., 2011).

Animals were treated according to French and European regulations of animals in research. SR’ authorization for use of animals in research is 91-116. There was no necessity of protocol authorization delivery by the Paris Centre-Sud Ethic committee for this work, as all experiments were performed on early 10hpf (hours post-fertilization) embryos, which are non-autonomous and have no nervous system yet.

### Embryo staging and fixation

Morphological criteria were taken to stage 10hpf and 10.5hpf embryos, in addition to their known developmental time. These included extent of blastopore closure (100% epiboly), the prominent tail bud and the lateral flat triangular shape of the ANP (Hinaux et al., 2011). Embryos were fixed in 4% paraformaldehyde in PBS, dehydrated in graded ethanol/PBS steps, and stored in methanol at −20°C.

### Whole-mount in situ hybridization (ISH)

ISH was carried out as previously described (Alié et al., 2018). Digoxigenin- and fluorescein-labeled riboprobes were prepared using PCR products as templates. cDNAs of interest were searched in our EST (expressed sequence tag) library (Hinaux et al., 2013). Clones in the library (pCMV SPORT6 vector) were: *rx3* (FO289986), *cxcr4b* (ARA0AAA96YA18), *pax6a* (ARA0AAA41YH22), *barhl2* (ARA0AEA6YL01), *nkx2*.1a (AY661435, (Menuet et al., 2007)), *zic1* (FO290256) (Devos et al., 2019), *lhx2* (EF175737) (Pottin et al., 2011), *emx3* (FO263072). For fluorescent *in situ* revelation, FITC- and Cy3-tyramides (*excitation* peaks at 491 and 555 nm and emission peaks at 516 and 569 nm, respectively) were prepared as described (Zhou and Vize, 2004). Embryos incubation in PBS containing 10uM DAPI and 1%DMSO 24 hours prior to dissection to stain nuclei and washed in PBS.

### Colorimetric ISH, image acquisition

Whole-mount embryos stained by colorimetric ISH were imaged on a Nikon AZ100 multizoom macroscope coupled to a Nikon digital sight DS-Ri1 camera, using the NIS software.

### Fluorescent ISH, neural plate dissections

Fluorescently labelled whole-mount embryos were transferred on Sylgard™-containing petri dish bathed in PBT (PBS + 0.3% Tween). Embryos were cut using forceps (Fine science Tools, #5 Ultra) in two pieces at the yolk equator. After yolk removal, green and/or red staining of the ANP served to dissect-out and isolate a piece of tissue containing the labeled ANP, under a fluorescent dissecting microscope. Samples were mounted in dorsal views in Vectashield (Vector) between glass slides and coverslips inside wells built with one reinforcement ring.

### Fluorescent image stack acquisition

Confocal image stacks were captured on a confocal Leica SP8 microscope using the Leica Application Suite software. Confocal image stacks were captured on a confocal Leica SP8 microscope using the Leica Application Suite software. Water immersion objective (HC FLUOTAR L 25x/0.95 W VISIR 2.50 Water) was used for all experiments except for “single cell” analyses which required x40 objective (HC PL APO 40x/1.10 W CORR CS2 0.65 Water 0.14–0.18) to improve resolution and hence better separate nuclei pixel intensities. Since we did not compare mean intensities between samples but relative fluorescent levels within each normalized sample, we choose the optimal laser power and photo-multiplication (PMT, ranging from 450 to 630) conditions for each sample acquisition. PMT gain compensation in the z axis was also applied to capture signals in ventral deeper regions when stacking the entire eyefield. Acquisitions were performed using sequential modes with optimization of emission wavelength range to avoid bleed-through of signals. Image stacks depth of entire neural plate typically ranged between 70 and 90 z steps depending on samples. Z step of 1 µm was used for each confocal acquisition. Image size and format were of 512×512 pixels and 8 bit, respectively.

### Neural plate staging: d factor

The anterior curvature of the ANP looks like a parabola (y=ax^2^), where (a) corresponds to the vertical scaling factor. For convenience of our experimental design we used the formula (y=1/dx2), where factor d is the inverse of a. Therefore, the higher d is, the flatter the curve of the ANP is, the less advanced convergence is. Conversely, a lower d factor indicates a strong curvature of the ANP, thus advanced tissue convergence. We generated a single image (512×512 pixels) containing a merge of nine parabolas ranging from d factors 1 to 9, in an orthonormal system (Figure 1). This allowed generating composite 2-channels images containing maximum intensity projection of single fluorescent ISH image stacks superposed to the parabolas grid, in which the ANR medial anterior point was positioned at (x=0, y=0). We then assessed to which parabola/ d factor, each sample fitted the best, by eye. Since ANP anterior curvatures were sometimes slightly asymmetric, we calculated the d factor for each sample as the mean of the two values estimated for the left and right side. We validated the accuracy of this method by merging images of similar d factors and controlling by eye shape correspondence. Overall, this method allowed accessing intrinsic neural temporality/stage for each sample, and comparing them according to this criterion.

### Morphometric analyses

#### 2D and 3D size analyses

Areas on colorimetric and fluorescence images were drawn by tracing contours manually and measured using FIJI (Schindelin et al., 2012).

Volumes on image stacks were calculated as follows: for each marker, the objects were segmented using Imaris (Bitplane) and a common method. For each channel, we created a corresponding cell using the default mode (settings: volume display). Cell body detection used the smooth option (Filter width 2um). Cell threshold (absolute intensity) did not split touching cells and was adapted for each sample. Finally, object properties were thereafter extracted (i.e. volumes). To compute the volume of the overlapping region between two markers, we used the “Surface-surface colocalization XTension” (Imaris). For Figure 5, we calculated volumes of the solo rx3+ zic1-expressing domain, by summing substracted surface areas of each plan from binarized image stacks.

For Figure 4E, we used the Measure stack plugin (FIJI) allowing the semi-automatic drawing of nkx2.1 domain contours in each z step, whose sum allowed calculation of volume of prospective hypothalamus.

For Figure 5DE and supplementary Figure 5, we apply the Make Binary plugin (FIJI) on image stacks, setting identical minimum and maximum pixel intensities for all samples compared. Symmetrical subtraction of binarized zic1 and rx3 stack channels using the Image Calculator plugin allowed generating zic1+rx3− and rx3+zic1-images. Segmented areas were calculated from maximum pixel intensity ventral half stack projections using the Wand Tool. Volume of the midline ventral rx3+zic1-domain was calculated by summing all z segmented areas from rx3 - zic1 subtracted images.

#### 2D and 3D renditions

We performed volume renditions using the 3D Viewer plugin (FIJI) and Imaris (see Figure legends).

### Image processing and analyses

Fluorescence signal measurements were performed on maximum intensity projections, whose contrasts were adjusted automatically, in order to optimize the dynamic range of pixel intensities. Background signals were subtracted and fluorescence levels then normalized by the maximal value measured. All fluorescence measurements were normalized.

### Plot profiles

We generated plot profiles from maximum intensity projections of 10 µm thick sections at a position close to mid-z-depth of the expression domains considered in the majority of the experiments. For Figure 4, 6 and supplementary Figure 5, maximum intensity projections corresponded to entire stacks. In Figure 4, we considered this variant for measures as it clearly showed the relative position of the two tissues, eyefield and prospective hypothalamus. For Figure 6, it reflected well the relative position of the pairs of markers considered in the bended prospective telencephalon. The regions of interest (ROI) selected for measurements are indicated in each figure. Due to inter-individual size differences, the length of each plot for rx3 (Figure 1), pax6/rx3 (Figure 3) and nkx2.1/rx3 (Figure 4) was normalized by the total distance, thus allowing inter-sample relative pixel intensity comparisons.

For specific analyses at domain boundaries (pax6/ barhl2 (Figure 2), emx3/zic1 and zic1/rx3 (Figure 7)), the ROI were 20 µm wide and 50 µm long line histograms.

For analyses in the D/V axis (Figure 3), we resliced image stacks entirely according to the indicated scheme to generate 3µm thick sagittal projections. We next quantified relative pixel intensities in D/V line histograms equally spaced in the A/P axis inside the rx3 domain.

### Single cell analyses

For individual cell fluorescence quantifications, we used 2 µm maximum intensity projection images. Nuclear staining (DAPI) served to segment “individual particles” in the ROI, allowing measurement of pixel intensity mean levels for all channels (single cells). As shown in supplementary Figure, fluorescence levels measured in individual nuclei areas reflected the fluorescence levels in the corresponding surrounding concentric zones, thus showing that measure of pixel intensity levels in the nuclear zones was a good proxy of individual of cell fluorescence levels. Again, as all fluorescence measurements were normalized, this allowed comparing relative fluorescent intensity levels – taken as a proxy of gene expression level – between cells within a given sample.

### Statistical analyses

p-values were calculated using the Mann-Whitney U test and Chi test in R, except from Figure 5A’’ (two-sided t-test with unequal variance). No statistical method was used to predetermine sample size. The experiments were not randomized and the investigators were not blinded to allocation during experiments and outcome assessment.

## Acknowledgements

We thank Stéphane Père and Krystel Saroul for taking care of our *Astyanax* colony. We warmly thank Romain Le Bars (RLB) at the I2BC Imaging Plateform facility for assistance on the Imaris and FIJI softwares. We also would like to thank Bruno Bozon and RLB for designing macros in FIJI.

Work supported by CNRS and grants from FRM (Fondation pour la Recherche Médicale DEQ2015030331745), UNADEV (Union Nationale des Aveugles et Déficients Visuels)/AVIESAN (Alliance nationale pour les sciences de la VIE et de la SANté) and RETINA France to SR. JTP received financial support from FRM and Becas Chile.

